# Distinctive Transcriptional and Microbial Signature in Cutaneous Acute Graft-vs-Host-Disease

**DOI:** 10.1101/2024.06.17.599323

**Authors:** Najla El Jurdi, Ashraf Shabaneh, Brittney Schultz, Owen Dean, Jinhua Wang, Shernan G. Holtan

**Affiliations:** Blood and Marrow Transplant Program, University of Minnesota; Cancer Bioinformatics, Masonic Cancer Center, University of Minnesota; Department of Dermatology, University of Minnesota; Department of Pediatrics, University of Minnesota

**Author notes:** **Correspondence:** Najla El Jurdi, MD, Assistant Professor of Medicine, Division of Hematology, Oncology and Transplantation, 420 Delaware Street SE, Minneapolis, MN 55455. **Data Availability:** Datasets will be publicly available on Gene Expression Omnibus (GEO) with accession number GSE263557. **Link to data specific repository page:** https://www.ncbi.nlm.nih.gov/geo/query/acc.cgi?acc=GSE263557. **Code Availability statement:** no additional custom code was generated for this analysis. ***Contribution:*** NEJ performed the bibliographic search and wrote the first version of the manuscript; NEJ, SGH contributed to the design; NEJ and BS performed the retrospective chart reviews; AS and JW performed the statistical analysis; all other authors edited and revised the manuscript.

**Keywords:** Genomics, microbiome data, GVHD, cutaneous

## Abstract

Skin acute graft-vs-host disease (aGVHD) is often first manifestation of GVHD, yet very few preclinical and clinical studies have focused on this target organ, leaving a critical information gap in the pathophysiology of GVHD. We hypothesized that analysis of host and microbiome gene expression could yield novel insights into the molecular and immunologic mechanisms underlying skin GVHD. Our objectives were to determine the differential host gene expression and microbiome profile of human skin aGVHD samples compared to normal skin, and aGVHD corticosteroid responders to non-responders. We performed RNA-Sequencing on lower arm biopsies from 45 patients compared to 10 healthy controls. Our findings suggest a distinctive transcriptional signature of cutaneous aGVHD, that could identify potentially actionable targets for prevention or treatment corticosteroid refractory disease. Our analysis suggests a key role of dendritic cells and macrophages, potentially mediated by differential expression of MIF, in the development of cutaneous aGVHD and corticosteroid responsiveness. Additionally, we describe a unique microbial signature in cutaneous aGVHD that includes skin microbes not previously described in this population.

## Introduction

Allogeneic hematopoietic cell transplantation (HCT) is frequently associated with significant toxicities, including graft-vs-host disease (GVHD) that increase morbidity and mortality of this lifesaving therapy. Mucosal damage, secondary to cytotoxic conditioning regimens, results in microbiome dysbiosis with altered antimicrobial peptide expression. This dysbiosis is one possible mechanism for stimulation of alloreactive T cell responses inciting GVHD. Decades of strong scientific evidence identified intestinal microbiota composition, diversity, and antimicrobial peptides as regulators of immune tolerance and allogeneic HCT outcomes^1,2^. However, the role of the skin microbiome in this critical patient population remains unexplored or clinically defined. Skin GVHD is the most common and often first manifestation of acute GVHD (aGVHD), a cutaneous manifestation of a systemic disease that can have an unpredictable treatment course. Skin, is the largest human organ, an immune organ and often the first line of defense against pathogens. Analysis of both host and microbiome gene expression could yield novel insights into mechanisms of response to corticosteroids. This study aims to explore molecular and immunologic mechanisms underlying skin GVHD, as well as the potential contributions of microbes to treatment outcomes. Our objectives were to determine simultaneously the differential host gene expression and microbiome profile of human skin GVHD samples compared to normal skin and within the aGVHD group to compare corticosteroid responders compared to non-responders.

## Methods

To maintain consistency and allow for comparative dual host and skin microbiome analysis, we performed RNA-Sequencing (RNA-seq) on 20 μm slices of formalin-fixed paraffin embedded (FFPE) diagnostic lower arm skin biopsies from 45 adult patients with histologic grade 2-2.5 acute GVHD (aGVHD). We compared these to 10 healthy controls from discarded surgical lower arm skin tissue. All skin samples were anatomically from the forearm. We assessed aGVHD treatment response at day 28 and identified 35 complete or partial responders (CR or PR) and 10 non-responders (NR) based upon their response to systemic immunosuppression with corticosteroids at day 28 of aGVHD therapy.

We extracted RNA from the FFPE biopsies using PureLink FFPE RNA Isolation Kit. FFPE samples yielded an adequate quantity of RNA for sequencing (median = 98717524.5 reads, range = 74316820 to 229178960). After extraction, we prepared the libraries with the SMARTer Stranded Total RNA Kit v2 and performed sequencing on a single lane of a NovaSeq S4 2×150-bp. We aligned and assembled sequences using a graph based alignment tool hisat2 and Cufflinks RNA-seq workflow, respectively; analyzed pathways using comprehensive gene set enrichment analysis web server Enrichr. For estimation of microbial RNA content, we analyzed reads that were not mapped to the human genome. Specifically, we used hisat alignment tool with option: -un-conc-gz to output unmapped pair ends as new FASTQ files; then we aligned unmapped pair ends using the hisat2 algorithm to the microbiome sequence database. We used PathSeq, a comprehensive pipeline tool to analyze microbial sequences, as a mapping and taxonomic classification algorithm to estimate abundance of candidate microbes using read counts assigned to each organism. We performed both unsupervised analyses as well as hypothesis-driven single gene and microbe analyses based upon targets of interest from the literature. This study was reviewed and approved by the University of Minnesota IRB.

## Data Records

Datasets will be publicly available on Gene Expression Omnibus (GEO) with accession number GSE263557 (https://www.ncbi.nlm.nih.gov/geo/query/acc.cgi?acc=GSE263557).

## Results and Discussion

### Patient and aGVHD Characteristics

Patients underwent HCT at the University of Minnesota between 2004 and 2015. A total of 45 patients were included, 58% were male, 24% received a matched related donor, 9% matched unrelated donor, 67% umbilical cord blood donor, and 53% underwent a reduced intensity conditioning. Median age at HCT was 36 years (interquartile range IQR, 15-52). Acute leukemia was the most common indication for transplant. GVHD prophylaxis regimen was predominantly cyclosporine with mycophenolate mofetil (82%). Median days to aGVHD onset was 34 days (IQR, 24-45) from the time of HCT. Maximus aGVHD grade was distributed as follows: Grade I 29%, Grade II 36%, Grade II 29%, and Grade IV 7%. All patients had skin aGVHD at diagnosis (clinically and at the time the diagnostic biopsy was obtained) with the majority of patients (73%) not having gastrointestinal manifestations of aGVHD.

### Skin aGVHD vs. normal control: Single Gene Differential Expression

We compared the differential expression of single genes and found immune and damage relevant genes were differentially expressed in aGVHD vs. normal skin controls. Specifically, 437 genes were at least 2-fold significantly differentially expressed in aGVHD vs. normal skin controls. **Table 2** shows the 10 most significantly differentially expressed genes. For example, PEA15 was 2.44 fold change higher in aGVHD and it is a gene encoding a death effector domain-containing protein that functions as a negative regulator of apoptosis that has a role implicated in drug resistant in breast and ovarian cancer as well as diabetes mellitus^3,4^.

**Table 1.**
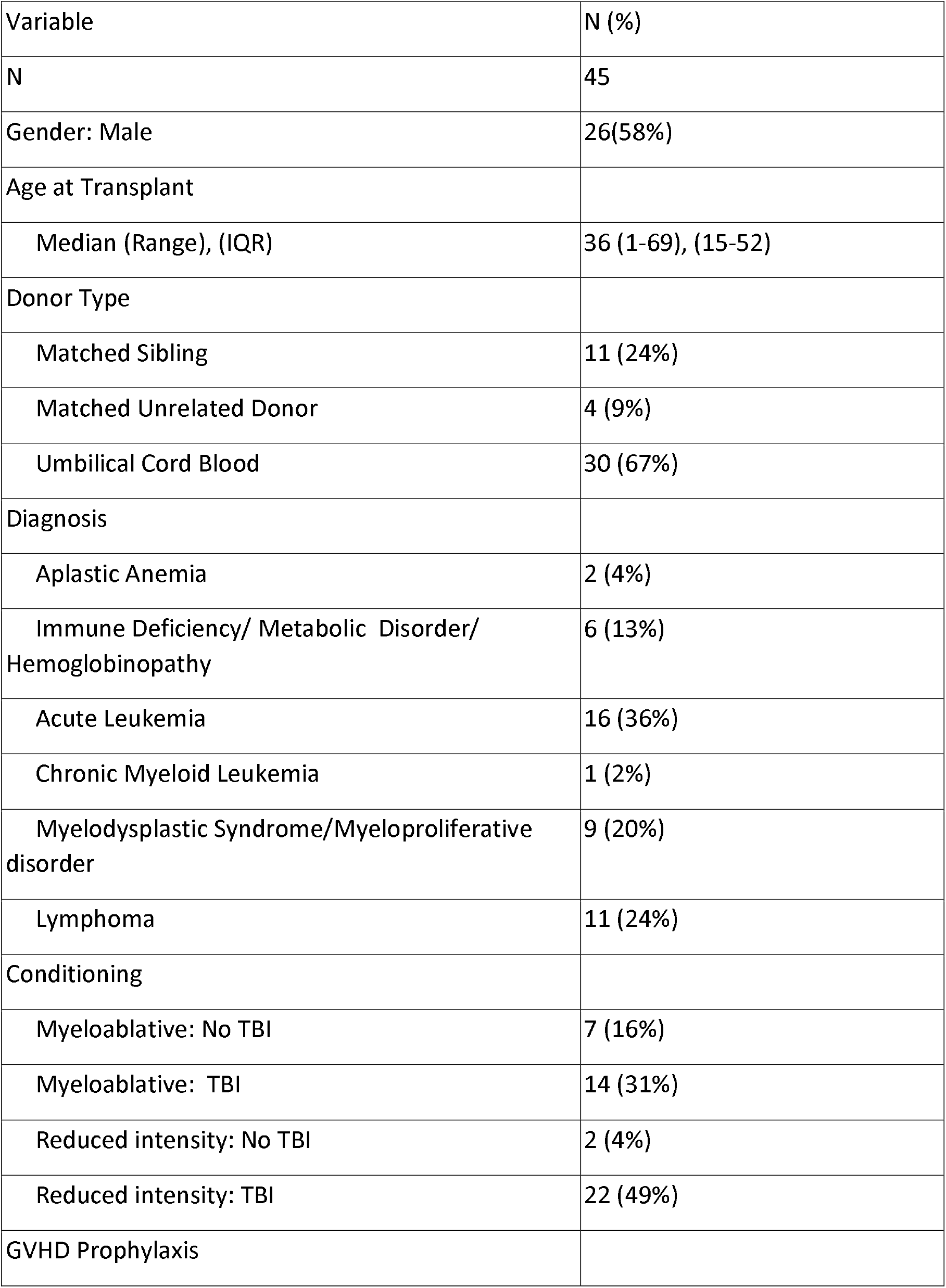
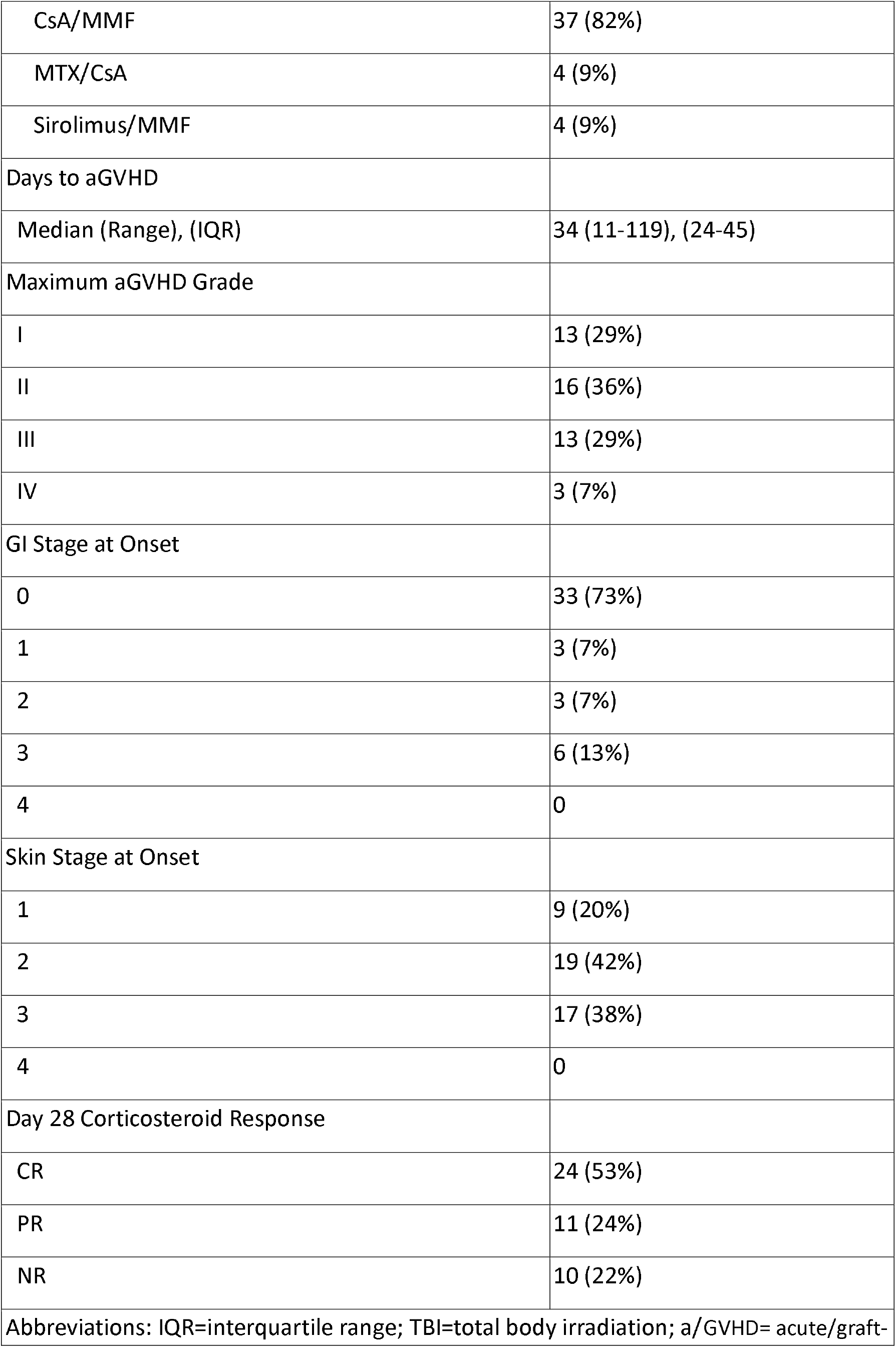
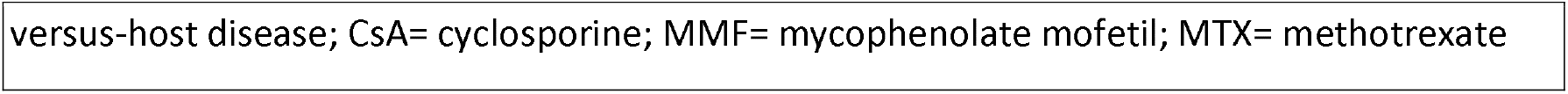
Patient, Disease, and GVHD characteristics.

**Table 2.**
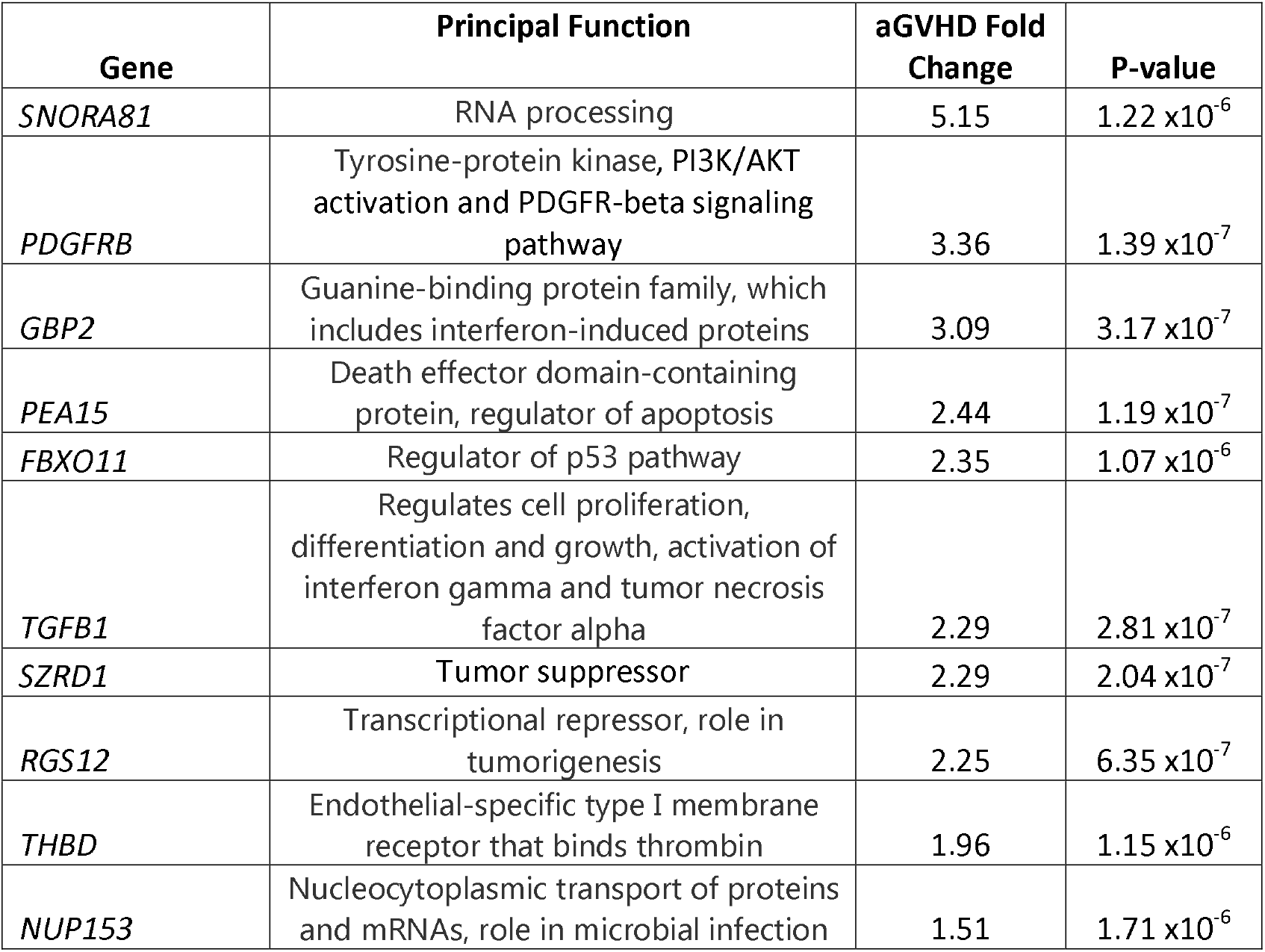
Single Gene Differential Expression of Skin aGVHD Compared to Normal Control.

| Gene | Principal Function | aGVHD Fold Change | P-value |
| --- | --- | --- | --- |
| *SNORA81* | RNA processing | 5.15 | 1.22 x10<sup>-6</sup> |
| *PDGFRB* | Tyrosine-protein kinase, PI3K/AKT activation and PDGFR-beta signaling pathway | 3.36 | 1.39 x10<sup>-7</sup> |
| *GBP2* | Guanine-binding protein family, which includes interferon-induced proteins | 3.09 | 3.17 x10<sup>-7</sup> |
| *PEA15* | Death effector domain-containing protein, regulator of apoptosis | 2.44 | 1.19 x10<sup>-7</sup> |
| *FBXO11* | Regulator of p53 pathway | 2.35 | 1.07 x10<sup>-6</sup> |
| *TGFB1* | Regulates cell proliferation, differentiation and growth, activation of interferon gamma and tumor necrosis factor alpha | 2.29 | 2.81 x10<sup>-7</sup> |
| *SZRD1* | Tumor suppressor | 2.29 | 2.04 x10<sup>-7</sup> |
| *RGS12* | Transcriptional repressor, role in tumorigenesis | 2.25 | 6.35 x10<sup>-7</sup> |
| *THBD* | Endothelial-specific type I membrane receptor that binds thrombin | 1.96 | 1.15 x10<sup>-6</sup> |
| *NUP153* | Nucleocytoplasmic transport of proteins and mRNAs, role in microbial infection | 1.51 | 1.71 x10<sup>-6</sup> |

### Skin aGVHD vs. control: Higher Expression of Interferon Responses and Inflammation Pathways

Enrichr Transcriptome analysis of those differentially expressed genes revealed a distinct pattern of pathway differentiation in aGVHD vs. controls. Immune and damage relevant pathways were enriched in aGVHD vs. controls, summarized in **Figure 1** showing the top 10 of 31 significantly enriched pathways, including: interferon alpha and gamma response, epithelial mesenchymal transition, IL-6/JAK/STAT3 signaling pathway, apoptosis, and p53, PI3K/mTOR signaling, IL-2/STAT5 pathways.

**Figure 1.**
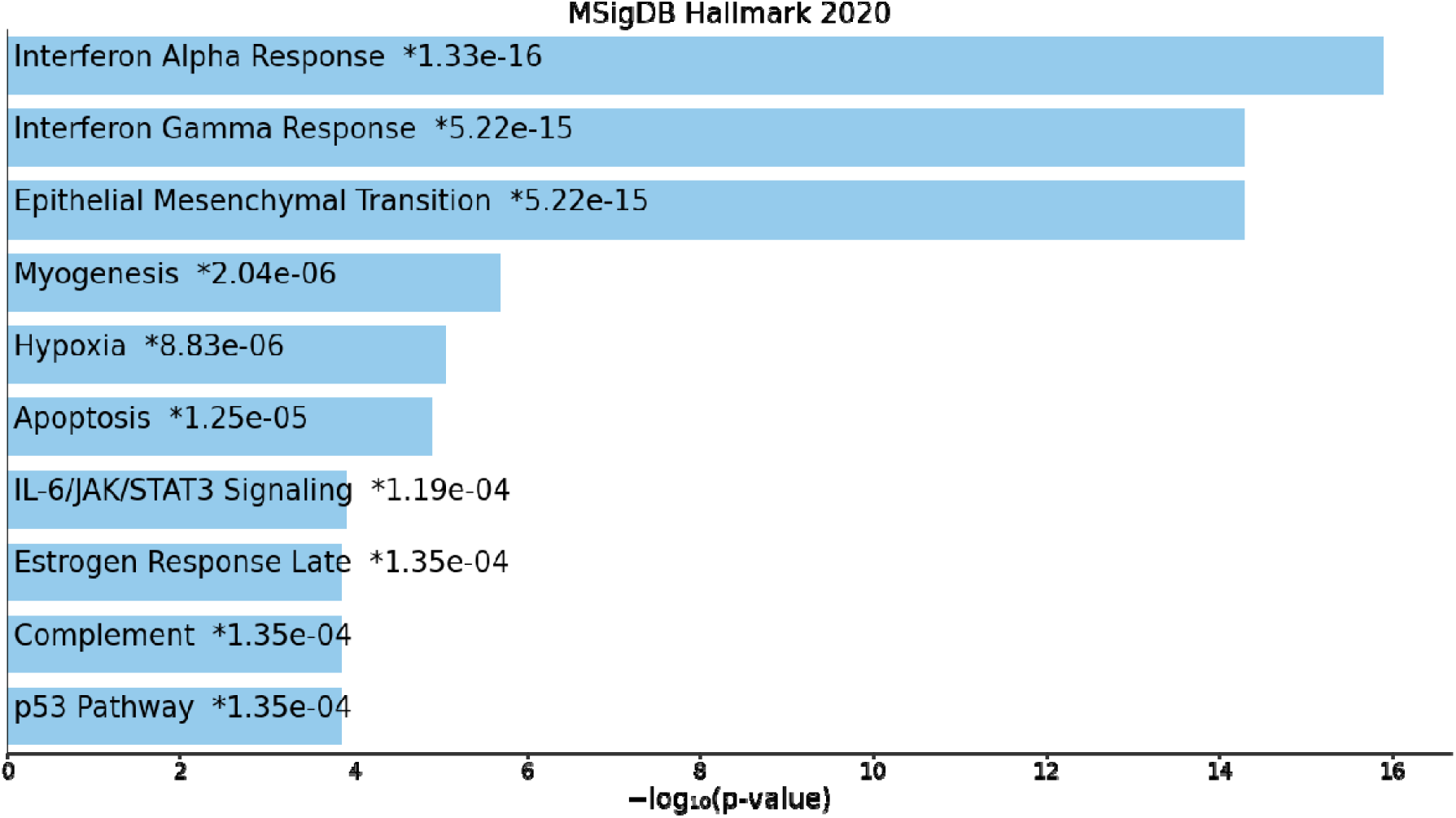
Enrichr analysis revealed a distinct pattern of pathway differentiation in aGVHD compared to controls.

### aGVHD Responder vs. Non-responder: Upregulation of MIF in Non-responders

We then grouped aGVHD samples by response to systemic immunosuppression with corticosteroids at day 28 of aGVHD therapy. We identified 35 complete or partial responders (CR/PR) and 10 non-responders (NR). Single gene analysis of the 2 groups revealed 5 genes that were significantly upregulated in responders (SRP14, AGTRAP, KATNBL1, C1QBP, and TMEM50A), and 3 genes that were significantly upregulated in non-responders (TEN1, RN7SL657F) including macrophage migratory inhibitory factor (MIF) a lymphokine involved in cell-mediated immunity, immunoregulation, and inflammation **(Figure 2)**.

**Figure 2.**
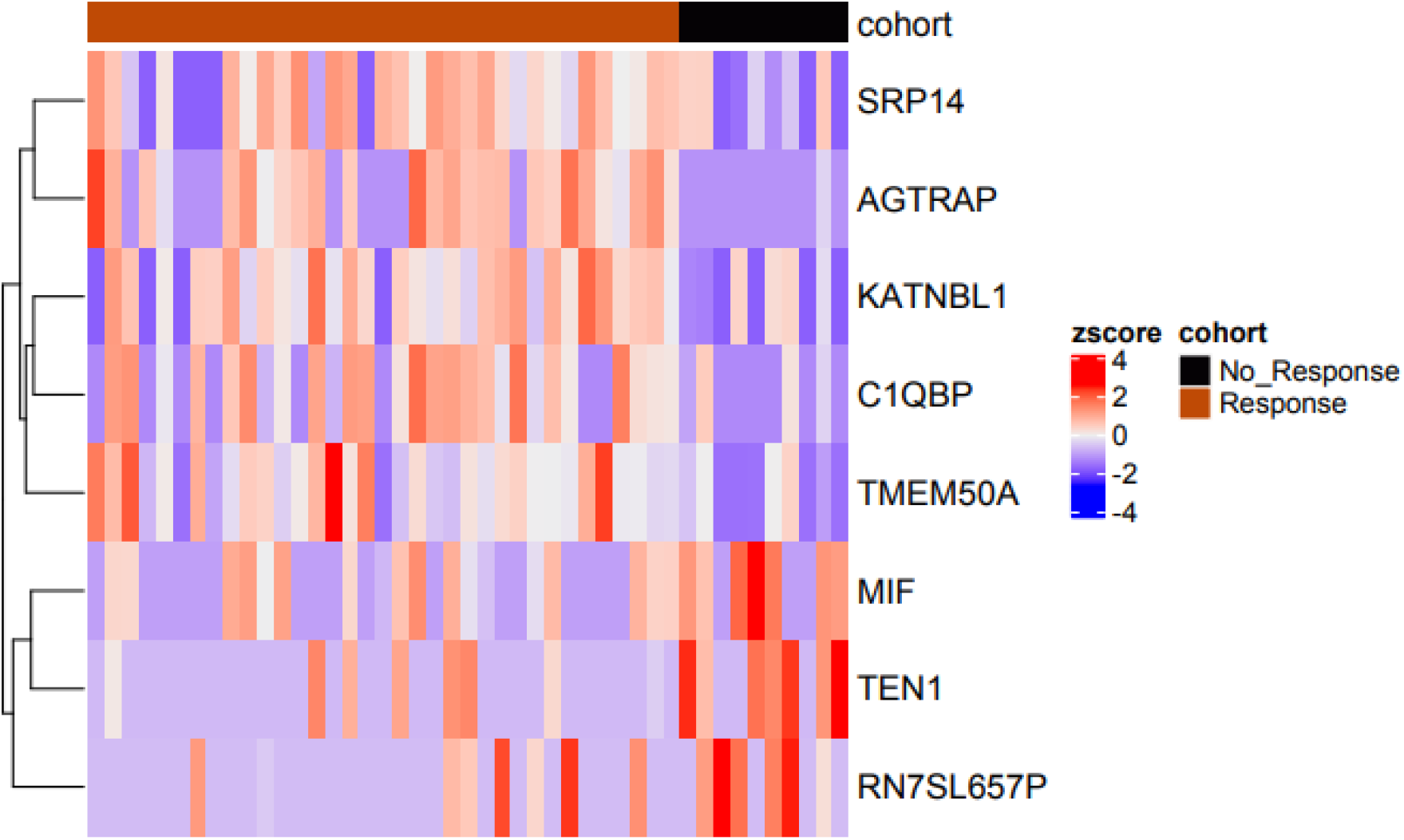
Heat Map of Single Gene Differential Expression of aGVHD Responders compared to Non-responders

MIF is a pro-inflammatory cytokine and is implicated in cancer with expression directly correlating with aggressive of certain subtypes^5^. Additionally, previous studies showed a significant increase in serum MIF levels (as well as skin and colonic MIF expression) in allogeneic HSCT recipients with aGVHD^6–9^. Interestingly, MIF plays a role in the regulation of macrophage function in host defense through the suppression of anti-inflammatory effects of glucocorticoids^10^ and seen here to be upregulated in non-responders, possibly explaining the lack of response to corticosteroids in this subgroup of patients.

### aGVHD Responder vs. Non-responder: Biologic Processes

**Figure 3a** summarizes the biologic processes upregulated in responders. We noted a higher expression of processes of ribosome assembly (cell growth), dendritic cell chemotaxis, and trophoblast migration (angiogenesis, tolerance) and negative regulation of pattern recognition-mediated inflammatory responses via retinoic acid-inducible gene I (RIG-I) and melanoma differentiation-associated gene 5 (MDA-5). RIG-I and MDA-5 function as a pattern recognition receptors in the RIG-1 pathways that are thought to have a key role in damage and pathogen-associated molecular patterns (DAMPs/PAMPs)-mediated inflammatory responses and stimulators for (host and donor-derived) antigen presenting cells (APCs)^11–13^. **Figure 3b** summarizes the biologic processes upregulated in non-responders. We noted an upregulation of processes involved in apoptosis, DNA damage, and cell aging along with negative regulation of macrophage migration and chemotaxis macrophage chemotaxis.

**Figure 3a.**
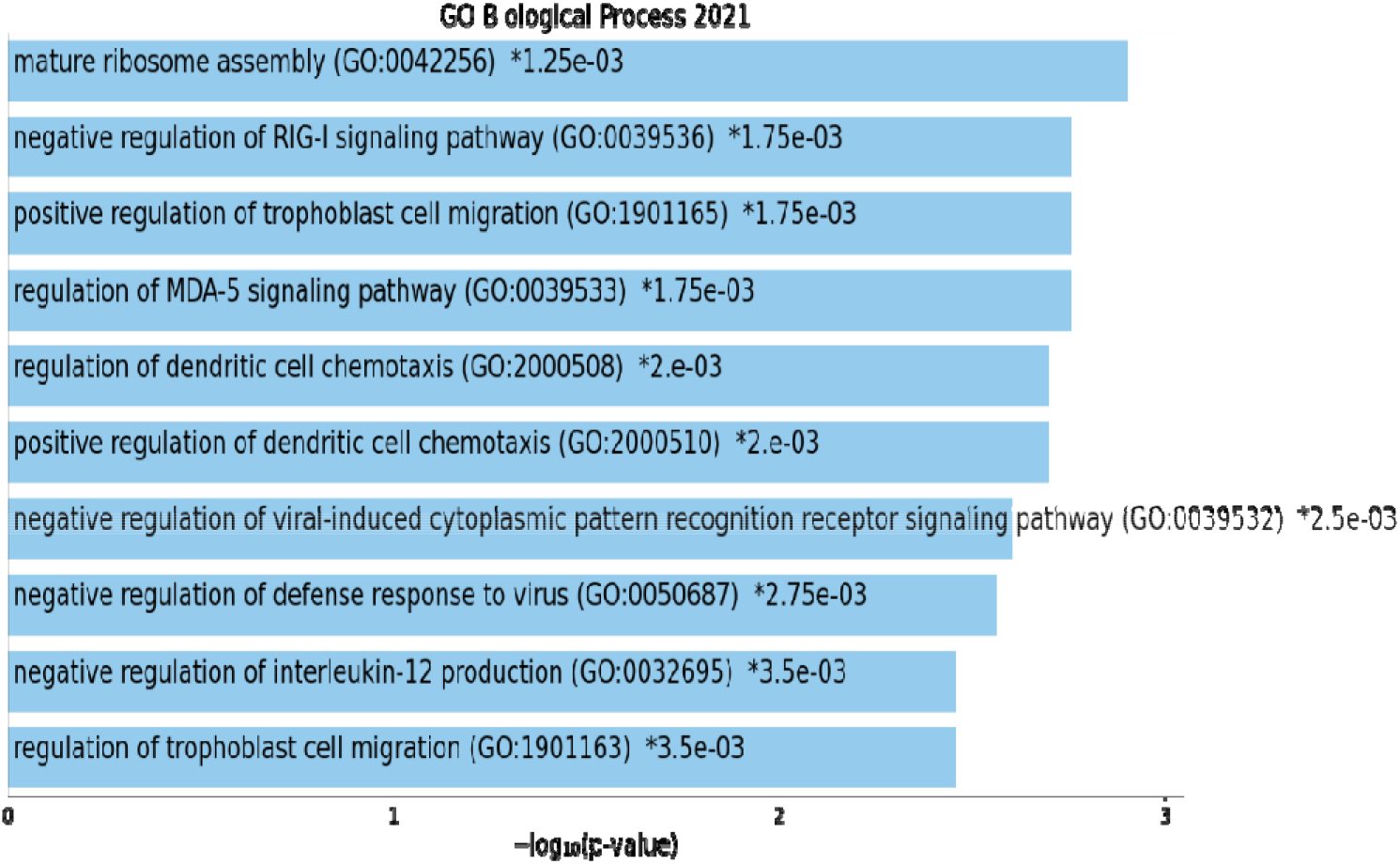
Biologic Processes Upregulated in aGVHD Responder compared Non-responder

**Figure 3b.**
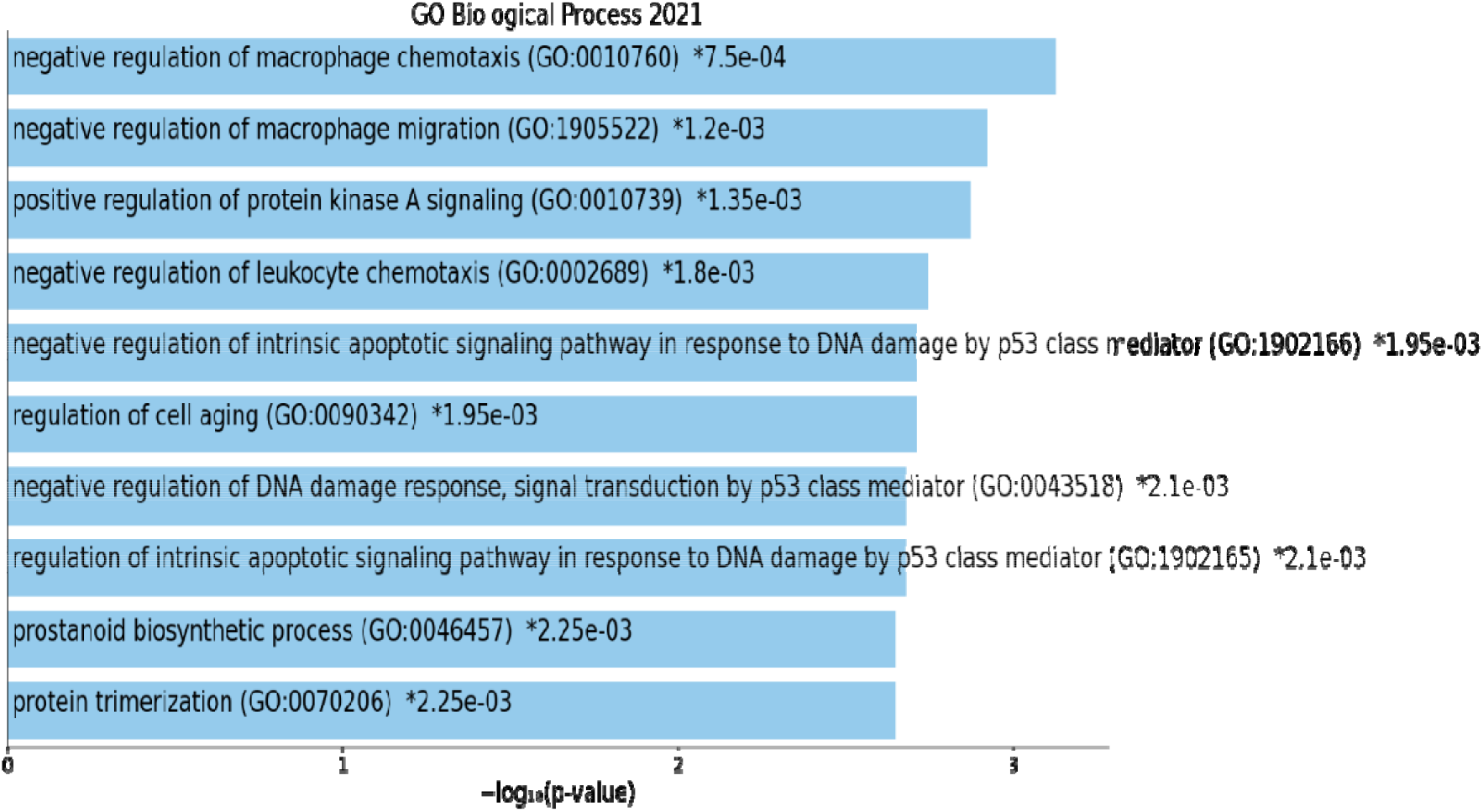
Biologic Processes Upregulated in Non-Responders compared to Responders

### CIBERSORT-Correlations Between Immune Cell Subsets

Using CIBERSORT to estimate cell composition and compare cell subsets across groups (control, non-responders and responders), we observed no statistically significant difference in any single cell type after adjustment for multiple comparisons. We then performed correlation analysis (Pearson’s r) and hierarchical clustering, to assess the presence and overall pattern of correlated immune cell types in estimated abundance. In comparison to the clustered heatmap in normal skin control, we noted a general loss of “normal” correlation in treatment responsive aGVHD samples, possibly relating to the cell-poor interface dermatitis that is a characteristic of cutaneous GVHD dermatitis^14^.

When examining the correlations between immune cell subsets in non-responsive aGVHD, we noted loss of Treg, high correlation of M0 macrophages with eosinophils and CD4 memory resting T-cells and gamma delta cells, as well as high correlation of M2 macrophages with CD8 T-cells.

### Microbiome Analysis

We then performed a preliminary analysis of the microbial differential abundance at the phylum level in aGVHD samples vs. controls. For estimation of microbial RNA content, we analyzed reads that were not mapped to the human genome and used PathSeq as a mapping and taxonomic classification algorithm to estimate abundance of candidate microbes. We notes a significant differential abundance at the phylum level in aGVHD samples vs. controls, not between treatment response groups (Table 3). Significant difference was in five phyla belonging to the kingdoms Eukaryota, Fungi and Archea, not Bacteria. On deeper hierarchical cluster analysis, we found a significant differential abundance of multiple taxa at the family, genus and species levels between aGVHD vs. controls and responders vs. non-responders. As shown in the heat map **Figure 4**, we noted an increased differential abundance of viruses in non-responders, decreased differential abundance of eukaryote in non-responders, and greater heterogeneity in differential abundance of bacteria between and within groups.

**Table 3.** Microbiome differential abundance at the phylum level in aGVHD samples vs. controls.

| Kingdom | Phylum | P- value | aGVHD vs. control Fold Change |
| --- | --- | --- | --- |
| Eukaryota | Phaeophyceae | 0.003 | 3.09 |
| Eukaryota | Euglenida | 0.02 | 2.80 |
| Fungi | Blastocladiomycota | 0.04 | 1.25 |
| Archaea | Thaumarchaeota | 0.04 | -0.77 |
| Fungi | Ascomycota | 0.02 | -0.80 |
| Archaea | Candidatus Korarchaeota | 0.03 | -1.47 |

**Figure 4.**
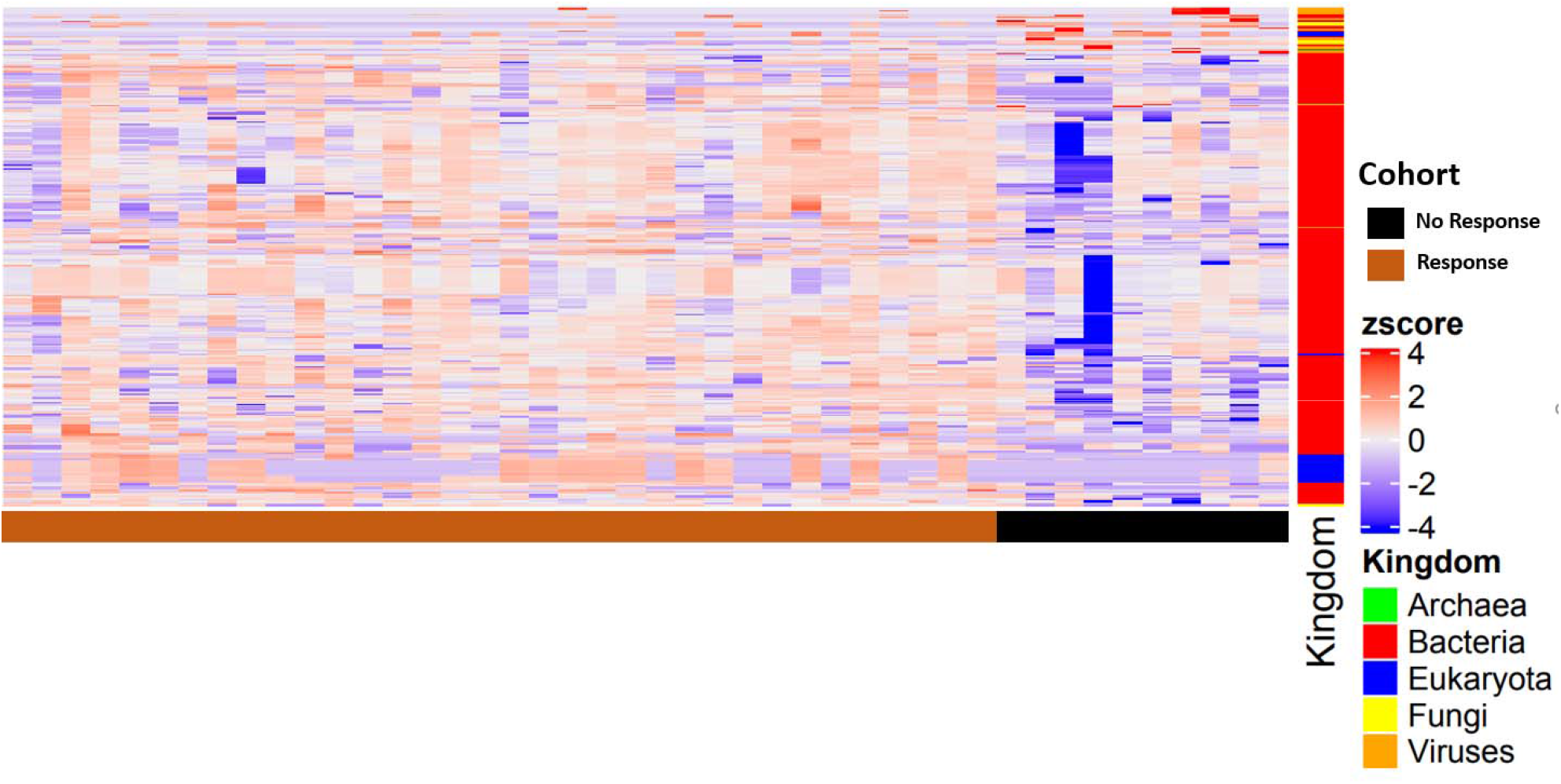
Heat map showing differential abundance of taxa at the family, genus and species levels between responders compared to non-responders

## Conclusions

Our findings show a distinctive transcriptional signature human cutaneous aGVHD, and further analysis and external validation could identify novel, potentially actionable, targets for prevention or treatment corticosteroid refractory disease. Our analysis suggests a key role of dendritic cells and macrophages, potentially mediated by differential expression of MIF, in the development of cutaneous aGVHD and corticosteroid responsiveness. Or data also suggests an important role of pattern recognition pathways, DNA damage, and cellular aging pathways that may facilitate human cutaneous GVHD development and disease severity. Additionally, we describe a unique microbial signature in cutaneous aGVHD that includes skin microbes not previously described in this population, extending beyond bacterial populations. This unique and rich dataset will be available to allow further investigations into mechanisms inciting cutaneous GVHD, affecting treatment sensitivity, and deciphering functional cutaneous microbial differences likely affecting local and systemic alloreactivity.

